# Enhanced bottom-up and reduced top-down fMRI activity is related to long-lasting non-reinforced behavioral change

**DOI:** 10.1101/363044

**Authors:** Rotem Botvinik-Nezer, Tom Salomon, Tom Schonberg

## Abstract

Behavioral change studies and interventions focus on self-control and external reinforcements as means to influence preferences. Cue-approach training (CAT) has been shown to induce preference changes lasting months following a mere association of items with a neutral cue and a speeded response, without external reinforcements. We utilized this paradigm to study preference representation and modification in the brain without external reinforcements. We scanned 36 participants with fMRI during a novel passive viewing task before, after and 30 days following CAT. We pre-registered the predictions that activity in regions related to memory, top-down attention and value processing underlie behavioral change. We found that bottom-up neural mechanisms, involving visual processing regions, were associated with immediate behavioral change, while reduced top-down parietal activity and enhanced hippocampal activity were related to the long-term change. Enhanced activity in value-related regions was found both immediately and in the long-term. Our findings suggest a novel neural mechanism of preference representation and modification. We suggest that non-reinforced change occurs initially in perceptual representation of items, which putatively lead to long-term changes in memory and top-down processes. These findings could lead to implementation of bottom-up instead of top-down targeted interventions to accomplish long-lasting behavioral change.

## Introduction

Changing behavior is key to solving a broad range of challenges in public health. Understanding how preferences are constructed and modified is a major challenge in the research of human behavior with broad implications, from basic science to offering long-lasting behavioral change programs (Marteau, Hollands, & Fletcher, 2012; Vlaev, Chater, Stewart, & Brown, 2011). Most behavioral interventions for treating conditions such as addictions and eating-disorders currently rely on reinforcements and effortful self-control (Wood & Neal, 2016). However, previous studies suggest that these interventions tend to fail in the long term (Christiansen, Bruun, Madsen, & Richelsen, 2007; Jeffery et al., 2000; Prochaska, Delucchi, & Hall, 2004).

In a recently introduced paradigm, named cue-approach training (CAT), preferences for snack food items were successfully modified in the absence of external reinforcements (Schonberg et al., 2014). In the CAT paradigm, the mere association of images of items with a cue and a speeded button-press response lead to preference changes lasting months following a single training session (Salomon et al., 2018; Schonberg et al., 2014). Current theories in the field of value-based decision-making would not predict that a simple association of an image with a neutral cue and button press will affect choices lasting months into the future. However, replicated results of over 30 samples in multiple laboratories show that participants significantly choose high-value paired items (*‘Go items’*) over high-value non-paired items *(‘NoGo items’*) following CAT (Bakkour et al., 2016, 2018; Bakkour, Lewis-Peacock, Poldrack, & Schonberg, 2017; Schonberg et al., 2014; Veling et al., 2017; Zoltak, Veling, Chen, & Holland, 2017). Salomon et al. (2018) recently showed that CAT can be used to change preferences towards various types of stimuli (snack food items, unfamiliar faces, fractal art images and positive affective images) with different types of cues (neutral auditory, aversive auditory and visual cues), demonstrating that the underlying mechanisms of the effect are general. Preference change following the task has been shown to last up to six months following a single training session lasting less than one hour, suggesting the task has potential to be translated into a real-world intervention and that it involves long-term memory components.

Training in the task is performed on single items and thus induces changes of preferences towards individual items, later manifested in the binary choice phase. The low-level nature of the task, involving neither external reinforcements nor high-level executive control, but rather sensory-motor associations, provides a unique opportunity to study in relative isolation preference representation and modification in the brain.

The underlying neural mechanisms driving this replicable long-lasting change remain largely unknown. Previous studies showed that eye-gaze during binary choices, following CAT, was drawn towards high-value Go items more compared to high-value NoGo items, even when the Go items were not chosen (Schonberg et al., 2014). Functional MRI (fMRI), during choices of high-value Go items alone and compared to choices of high-value NoGo items, demonstrated an amplified BOLD signal in the ventro-medial prefrontal cortex (vmPFC) (Schonberg et al., 2014), a region associated with value-based decision-making (Chib, Rangel, Shimojo, & O’Doherty, 2009). Together, these results indicate the involvement of attentional mechanisms and a neural signature of the value change during choices of Go compared to choices of NoGo items. Overall, these previous studies demonstrated that fMRI data during training and choices were not sufficient to reveal the underlying neural mechanisms of preference change induced by the task via training of individual items in the absence of external reinforcements.

Therefore, here we aimed to study how preferences toward individual items are changed in the task and uncover how individual items’ value is represented and modified in the brain even without external reinforcements. To do so, we introduce a novel passive viewing task, whereby pictures of snack food items are individually presented on the screen, while participants perform a sham counting task. This task was performed and scanned before, after and one month following CAT. By comparing fMRI activity during this task, we aimed to test the different neural responses to the same images of Go versus NoGo items after training compared to baseline, as well as for the first time the neural changes one month following training. Regions in the brain showing preference-related functional plasticity immediately after training and one month later, could reveal a novel mechanism of preference representation in the brain and specifically indicate how non-externally reinforced training leads to robust long-lasting preference changes.

We hypothesized that preference changes are dependent on attentional and memory-related mechanisms, affecting value representation. Based on previous findings (Schonberg et al., 2014; Veling et al., 2017), we hypothesized that attention-related processes are involved in the behavioral change following CAT and focused our predictions on top-down attention-related regions. Moreover, we hypothesized that memory processes are involved in the neural mechanism underlying the behavioral change following CAT, in the short- and long-term. This is following the findings that the preference changes induced by the task lasted for months after a single training session (Salomon et al., 2018; Schonberg et al., 2014) and based on recent theories for the involvement of memory in value-based decision-making (Shadlen & Shohamy, 2016; Shohamy & Daw, 2015; Weber & Johnson, 2006; Wimmer & Shohamy, 2012). Finally, as was previously demonstrated during the choice phase of the cue-approach task (Bakkour et al., 2017; Schonberg et al., 2014), we hypothesized we will observe neural changes in pre-frontal value-related regions. In our pre-registered hypotheses (https://osf.io/q8yct/), we predicted greater BOLD activity after CAT in response to high-value Go items in episodic memory-related regions in the medial temporal lobe (Brown, Staresina, & Wagner, 2015), top-down attention-related dorsal parietal cortex (Cabeza, Ciaramelli, Olson, & Moscovitch, 2008; Corbetta & Shulman, 2002) and prefrontal value-related regions (Kable & Glimcher, 2009; Padoa-Schioppa, 2011). In addition, we hypothesized we will replicate previous CAT results showing a significant behavioral effect of choosing high-value Go over high-value NoGo items during the binary choice probe phase and enhanced BOLD activity in the vmPFC during choices of high-value Go items (Bakkour et al., 2016, 2018, 2017; Salomon et al., 2018; Schonberg et al., 2014; Veling et al., 2017; Zoltak et al., 2017).

Understanding the neural mechanisms underlying non-reinforced behavioral change could potentially set the ground for new theories of value-based decision-making, and for new behavioral change interventions targeting automatic processes for long-lasting change, benefiting the lives of millions around the world.

## Materials and Methods

### Data sharing

We pre-registered our sample size, hypotheses and analyses plan (prior to final full analyses) on the Open Science Framework (OSF; https://osf.io/x6hsq/). The behavioral data and analysis codes are also available on the pre-registered OSF project. Imaging data are available in Brain Imaging Data Structure (BIDS) format (Gorgolewski et al., 2016) on OpenNeuro (as well as FSL’s design.fsf files, confounds.tsv files and the SVC masks): https://openneuro.org/datasets/ds001417. Unthresholded and thresholded statistical images of the imaging results are available on NeuroVault (Gorgolewski et al., 2015): https://neurovault.org/collections/TTZTGQNU/.

### Participants

Forty healthy right-handed participants took part in this experiment. The sample size was chosen before data collection and pre-registered during data collection (https://osf.io/kxh9y/). We initially planned to collect n = 35 participants based on a power-analysis using previous imaging CAT sample (Schonberg et al., 2014), aimed to detect with 80% power significant (p = .05) activity in the vmPFC. During data collection, we found that attrition rates were higher than expected, thus the planned sample size was increased to n = 40 (before exclusions and attrition), and re-registered. The total number of valid participants included in the final analyses of the first session is n = 36 (19 females, age: mean = 26.11, SD = 3.46 years). Of these participants, n = 27 returned for an additional one-month follow-up session (15 females, age: mean = 26.15, SD = 3.44 years).

All participants had normal or corrected-to-normal vision and hearing, no history of eating disorders or psychiatric, neurologic or metabolic diagnoses, had no food restrictions and were not taking any medications that would interfere with the experiment. Participants were asked to refrain from eating for four hours prior to arrival to the laboratory (Schonberg et al., 2014). All participants gave informed consent. The study was approved by the institutional review board at the Sheba Tel Hashomer Medical Center and the ethics committee at Tel Aviv University.

#### Exclusions

A total of four out of the 40 participants were excluded: One participant due to incompletion of the experiment, one based on training exclusion criteria (7.5% false alarm rate during training) and two participants with incidental brain findings; resulting in n = 36 valid participants.

### Stimuli

Sixty color images of familiar local snack food items were used in the current experiment. Images depicted the snack package and the snack itself on a homogenous black rectangle sized 576 x 432 pixels (see Supplementary Table 1; Stimuli dataset was created in our lab and is available online at http://schonberglab.tau.ac.il/resources/snack-food-image-database/). All snack food items were also available for actual consumption at the end of the experiment. Participants were presented with the real food items at the beginning of the experiment in order to promote incentive compatible behavior throughout the following tasks.

### Experimental procedures

The general task procedure was similar to previous studies with CAT (Salomon et al., 2018; Schonberg et al., 2014), and is presented in Figure 1. In order to test for functional changes in the neural response to the individual items following CAT, we added a new passive viewing task before, after and one month following training.

**Figure 1.**
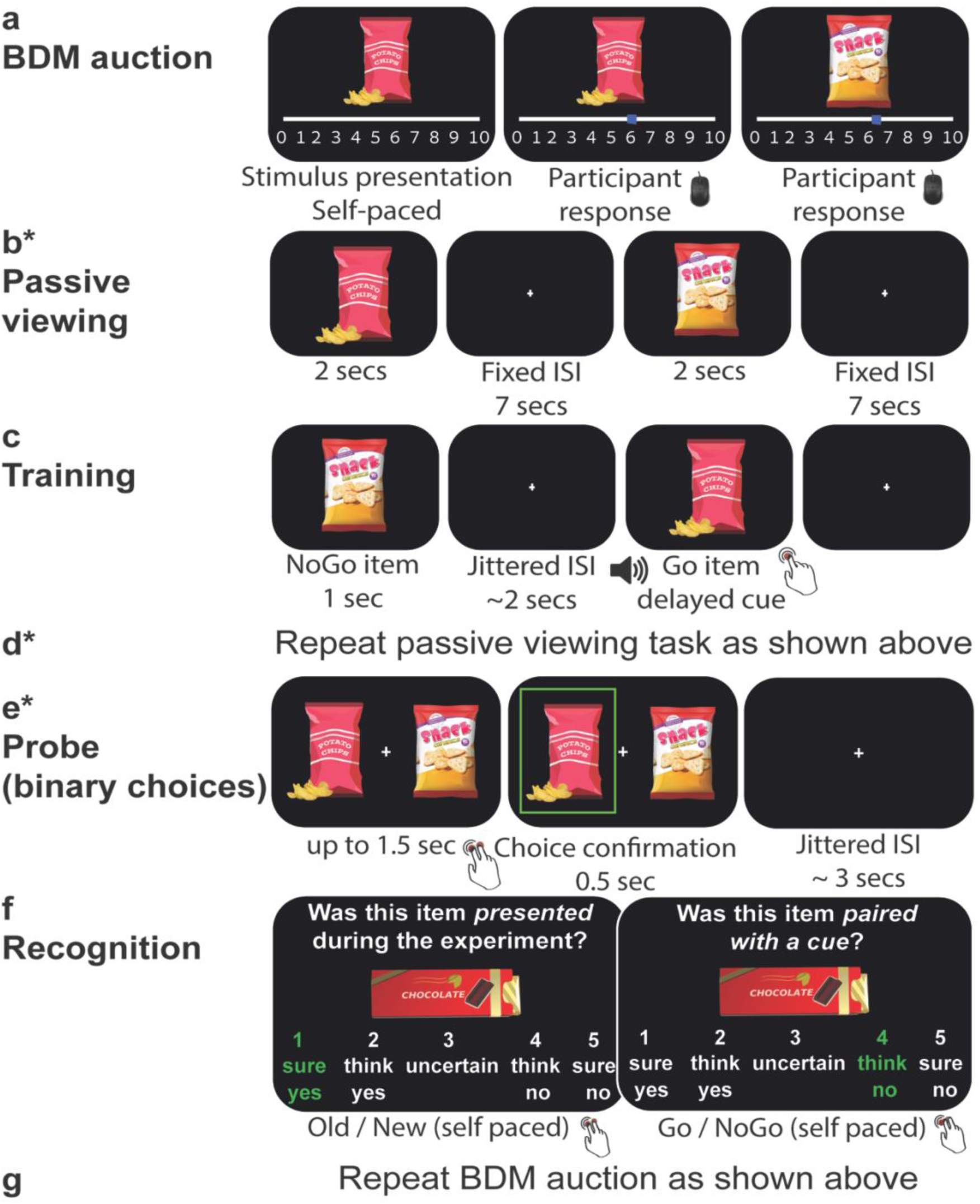
Outline of the experimental procedures: *Procedures performed inside the MRI scanner are marked with an asterisk.* ***(a)*** *Initial preferences were evaluated using the Becker–DeGroot–Marschak (BDM) auction procedure.* ***(b)*** *In the passive viewing task, items are individually presented on the screen.* ***(c)*** *Cue-approach training: Participants were instructed to press a button as fast as they could whenever they heard an auditory cue. Go items were consistently paired with the cue and button press response, while NoGo items were not.* ***(d)*** *The passive viewing task was repeated after training.* ***(e)*** *In the probe task, participants chose their preferred item between pairs of items with similar initial subjective preferences, one Go and one NoGo item.* ***(f)*** *A recognition memory task.* ***(g)*** *The BDM auction was repeated. Tasks d-g were performed again in the one-month follow-up session.*

First, we obtained the subjective willingness to pay (WTP) of each participant for each of the 60 snack food items using the Becker-DeGroot-Marschak (BDM) auction procedure (Becker, DeGroot, & Marschak, 1964), performed outside the MRI scanner (see Figure 1a,g). Then, participants entered the scanner and completed two passive viewing runs while scanned with fMRI (see Figure 1b,d), followed by anatomical and diffusion-weighted imaging (DWI) scans. Afterwards, participants went out of the scanner and completed cue-approach training (CAT) in a behavioral testing room at the imaging center (see Figure 1c). They then returned to the scanner and were scanned again with anatomical and DWI. Then, they were scanned with fMRI while performing two more runs of the passive viewing task and four runs of the probe phase, during which they chose between pairs of items (see Figure 1e). Finally, participants completed a recognition task outside the scanner (see Figure 1f). As the last task of the first session, they again completed the BDM auction to obtain their WTP for the snacks.

Approximately one month after the first session of the experiment, participants returned to the lab. They entered the scanner, were scanned with anatomical and DWI scans and completed two passive viewing runs as well as another probe phase (without additional training). Finally, participants completed the recognition and BDM auction parts, outside the scanner.

Anatomical and diffusion-weighted imaging data were obtained for each participant before, immediately after and one month following training. Analyses of diffusion data are beyond the scope of this paper.

#### Initial preferences evaluation (see Figure 1a,g)

In order to obtain initial subjective preferences, participants completed the BDM auction procedure (Becker et al., 1964). Participants first received 10 Israeli Shekels (ILS; equivalent to ∼2.7$ US). During the auction, 60 snack food items were presented on the screen one after the other in random order. For each item, participants were asked to indicate their willingness to pay (WTP) for the presented item. Participants placed their bid for each item using the mouse cursor along a visual analog scale, ranging from 0-10 ILS (task was self-paced). Participants were told in advance that at the end of the experiment, the computer will randomly generate a counter bid ranging between 0 - 10 ILS (with 0.5 increments) for one of the sixty items. If the bid placed by the participant exceeds the computer’s bid, she or he will be required to buy the item for the computer’s lower bid price. Otherwise, the participant will not be allowed to buy the snack but gets to retain the allocated 10 ILS. Participants were told that at the end of the experiment, they will stay in the room for 30 minutes and the only food they will be allowed to eat is the snack (in case they “won” the auction and purchased it). Participants were explicitly instructed that the best strategy for this task was to indicate their actual WTP for each item.

#### Item selection

For each participant, items were rank ordered from 1 (highest value) to 60 (lowest value) based on their WTP. Then, 12 items (ranked 7-18) were defined as high-valued items to be used in probe, and 12 items (ranked 43-54) were defined as low-valued items to be used in probe. Each group of twelve items (high-value or low-value) was split to two sub groups with identical mean rank. Six of the 12 items were chosen to be paired with the cue during training (Go items; training procedures are described in the following sections), and the other six were not paired with the cue during training (NoGo items). This allowed us to pair high-value Go and high-value NoGo items, or low-value Go and low-value NoGo items, with similar initial WTPs, for the probe binary choices. To maintain 30% frequency of Go items during training, similar to previous studies with CAT (Bakkour et al., 2016, 2017; Salomon et al., 2018; Schonberg et al., 2014), we used 16 additional NoGo items that were also used during training and passive viewing (40 snacks overall; see Supplementary Figure 1 for a detailed description of all stimuli allocation).

#### Passive viewing task (see Figure 1b,d)

The task was performed inside the scanner, while participants were scanned with fMRI. This new task was introduced to evaluate the functional changes in the neural response to the individual items following CAT. The neural signature of the participants’ response to each of the individual items was obtained in three different time points: A baseline measurement before CAT, after CAT and in a one-month follow-up. In this task, participants passively viewed a subset of 40 items, which were also presented during training (see item selection section and Supplementary Figure 1). The task consisted of two runs (in each session). On each run, each of the 40 items was presented on the screen for a fixed duration of two seconds, followed by a fixed inter-stimulus interval (ISI) of seven seconds. Items were presented in random order. To ensure participants were observing and processing the presented images, we asked them to perform a sham task of silently counting how many items were of snacks containing in a new package either one piece (e.g. a ‘Mars’ chocolate bar) or several pieces (e.g. a ‘M&M’ snack). At the end of each run, participants were asked how many items they counted. Task instructions (count one / several) were counterbalanced between runs for each participant. The time elapsed between the two runs before training and two runs after training was about two hours (including cue-approach training, anatomical and diffusion weighted scans before and after training and time to exit and enter the scanner).

#### Cue-approach training (see Figure 1c)

Training was performed outside the scanner. The training task included the same 40 items presented in the passive viewing task. Each image was presented on the screen one at a time for a fixed duration of one second. Participants were instructed to press a button on the keyboard as fast as they could when they heard an auditory cue, which was consistently paired with 30% of the items (Go items). Participants were not informed in advance that some of the items will be consistently paired with the cue, or the identity of the Go items. The auditory cue consisted of a 180ms-long sinus wave function. The auditory cue was heard initially 750ms after stimulus onset (Go-signal delay, GSD). To ensure a success rate of around 75% in pressing the button before stimulus offset, we used a ladder technique to update the GSD. The GSD was increased by 16.67ms following every successful trial and decreased by 50ms if the participant did not press the button or pressed it after the offset of the stimulus (1:3 ratio). Items were followed by a fixation cross that appeared on the screen for a jittered ISI with an average duration of two seconds (range 1-6 seconds). Each participant completed 20 repetitions of training, each repetition included all 40 items presented in a random order. A short break was given following every two training repetitions, in which the participants were asked to press a button when they were ready to proceed. The entire training session lasted about 40-45 minutes, depending on the duration of the breaks, which were controlled by the participants.

#### Probe (see Figure 1e)

Probe was conducted while participants were scanned with fMRI. The probe phase was aimed to test participants’ preferences following training. Participants were presented with pairs of items that had similar initial rankings (high-value or low-value), but only one of the items in each pair was associated with the cue during training (e.g. high-value Go vs. high-value NoGo). They were given 1.5 seconds to choose the item they preferred on each trial, by pressing one of two buttons on an MRI-compatible response box. Their choice was highlighted for 0.5 second with a green rectangle around the chosen items. If they did not respond on time, a message appeared on the screen, asking them to respond faster. A fixation cross appeared at the center of the screen between the two items during each trial, as well as during the ISI, which lasted on average three seconds (range 1-12 seconds).

The probe phase consisted of two blocks. On each block, each of the six high-value Go items were compared with each of the six high-value NoGo items (36 comparisons), as well as each of the six low-value Go items with each of the six low-value NoGo items. Thus, overall there were 72 pairs of Go-NoGo comparisons (each repeated twice during probe, once on each block). In addition, on each block we compared each of two high-value NoGo items versus each of two low-value NoGo items, resulting in four probe pairs that were used as “sanity checks” to ensure participants chose the items they preferred according to the initial WTP values obtained during the BDM auction. Each probe block was divided to two runs, each consisted of half of the total 76 unique pairs (38 trials on each run). All pairs within each run were presented in a random order, and the location of the items (left/right) was also randomly chosen. Choices during the probe phase were made for consumption to ensure they were incentive-compatible. Participants were told that a single trial will be randomly chosen at the end of the experiment and that they will receive the item they chose on that specific trial. The participants were shown the snack box with all snacks prior to the beginning of the experiment.

#### Recognition task (see Figure 1f)

Participants completed a recognition task, outside the scanner. In this task, the items from the probe phase, as well as an equal number of new items, were presented on the screen one by one and they were asked to indicate for each item whether or not it was presented during the experiment and whether or not it was paired with the cue during training. The first five participants completed a binary version of the recognition task: They first completed the old/new recognition task for all the items, and were then presented again with all the items they recognized as old items and were asked whether or not each item was paired with the cue (Go / NoGo recognition task). In the follow-up session, they were again presented with the items they indicated were old items (in Session 1), in a random order, and were asked again whether each item was paired with the auditory cue or not. In this version, each response was a binary yes/no response (“Was this item presented during the experiment?”). The rest of the participants completed a different version of the task. For each answer, they had five possible responses: Certain Yes, Think Yes, Uncertain, Think No or Certain No. The items were presented one by one on the screen, and for each item the participant was first asked whether this item was presented during the experiment and then, independent of the response to the first question, was asked whether or not it was paired with a cue during training.

#### One-month follow–up session

All participants were invited to the follow-up session approximately one month after training. A subset of 27 participants returned to the lab and completed the follow-up session. They were scanned with anatomical and diffusion protocols, completed two passive viewing runs and performed another probe while scanned with fMRI protocols, similar to the first session. In the follow-up session, the probe included the same pairs as the probe of the first session, presented in a new random order. Afterwards, participants completed another session of the recognition task and a third BDM auction, both outside the scanner in the testing room.

### Behavioral analysis

#### Probe

Similar to previous studies using cue-approach task (Salomon et al., 2018; Schonberg et al., 2014), we performed a repeated-measures logistic regression to compare the odds of choosing Go items against chance level (log-odds = 0; odds ratio = 1) for each trial type (high-value / low-value). We also compared the ratio of choosing the Go items between high-value and low-value pairs. These analyses were conducted for each session separately.

#### Recognition

In order to similarly analyze the recognition data across the two versions of the task, responses from the second version (i.e. with the confidence levels) were converted to binary yes/no answers. “Uncertain” was considered a wrong answer. In order to test whether Go items were better remembered than NoGo items following CAT, we compared the hit rate (the percent of old items that were correctly recognized as such) as well as response time (RT) in the old/new recognition task between Go and NoGo items that were included in the Go versus NoGo probe pairs (i.e. six high-value Go items, six high-value NoGo items, six low-value Go items and six low-value NoGo items). For the RT analysis, we only included trials with correct responses, under the assumption that shorter RTs for correct responses reflect better memory, while shorter RTs for incorrect responses do not.

It should be noted that this task was completed immediately after the probe task, both in the first and in the follow-up session. Hence, in the follow-up session the old/new task again tested memory for the same session, rather than long-term memory of the first session. Therefore, the Go / NoGo recognition task may be a better indication for long-term memory, but for the associations created during training (between the cue and Go items) rather than for the items themselves. Moreover, in the binary version of the task during the follow-up session (the first four participants in the follow-up session), only the Go/NoGo recognition task was performed. In addition, the recognition task (both versions and sessions) was self-paced, and thus the RT measure included outliers, for example when participants took a break; Therefore, RTs longer than three standard deviations above the mean for each version of the task were excluded from analysis.

#### Auction

Similar to the original CAT study (Schonberg et al., 2014), we tested whether WTP was changed over time differently for Go versus NoGo items. We computed ΔWTP (WTP after minus WTP before) for each item and each participant. Then, we used a repeated-measures linear regression with the ΔWTP as dependent variable and item type (Go / NoGo) and WTP before as independent variables. We were interested in the main effect of item type, i.e. whether ΔWTP was different for Go versus NoGo items. This analysis was performed separately for high-value and low-value items, as well as for the short-term change (after CAT minus before CAT) and the long-term change (one month following CAT minus before CAT).

### MRI data acquisition

Imaging data were acquired using a 3T Siemens Prisma MRI scanner with a 64-channel head coil, at the Strauss imaging center on the campus of Tel Aviv University. Functional data were acquired using a T2*-weighted echo planer imaging sequence. Repetition time (TR) = 2000 ms, echo time (TE) = 30 ms, flip angle (FA) = 90°, field of view (FOV) = 224 × 224 mm, acquisition matrix of 112 × 112. We positioned 58 oblique axial slices with a 2 × 2 mm in plane resolution 15° off the anterior commissure-posterior commissure line to reduce the frontal signal dropout (Deichmann, Gottfried, Hutton, & Turner, 2003), with a space of 2 mm and a gap of 0.5 mm to cover the entire brain. We used a multiband sequence (Moeller et al., 2010) with acceleration factor = 2 and parallel imaging factor (iPAT) = 2, in an interleaved fashion. Each of the passive viewing runs consisted of 180 volumes and each of the probe runs consisted of 100 volumes. In addition, in each scanning session (before, after and one month following training) we acquired high-resolution T1w structural images using a magnetization prepared rapid gradient echo (MPRAGE) pulse sequence (TR = 1.75 s, TE = 2.59 ms, FA = 8°, FOV = 224 × 224 × 208 mm, resolution = 1 × 1 × 1 mm for the first five participants; TR = 2.53 s, TE = 2.88 ms, FA = 7°, FOV = 224 × 224 × 208 mm, resolution = 1 × 1 × 1 mm for the rest of the sample. Protocol was changed to enhance the T1w contrast and improve registration of the functional data to the standard space).

### fMRI preprocessing

Raw imaging data in DICOM format were converted to NIfTI format with dcm2nii tool (Li, Morgan, Ashburner, Smith, & Rorden, 2016). The NIfTI files were organized according to the Brain Imaging Data Structure (BIDS) format v1.0.1 (Gorgolewski et al., 2016). These data are publicly shared on OpenNeuro (https://openneuro.org/datasets/ds001417). Preprocessing of the functional imaging data was performed using fMRIprep version 1.0.0-rc8 (Esteban et al., 2019), a Nipype based tool. Each T1 weighted volume was corrected for bias field using N4BiasFieldCorrection v2.1.0 and skull stripped using antsBrainExtraction.sh v2.1.0 (using OASIS template). Cortical surface was estimated using FreeSurfer v6.0.0 (Dale, Fischl, & Sereno, 1999). The skull stripped T1 weighted volumes were co-registered to skull stripped ICBM 152 Nonlinear template version 2009c (Fonov VS, Evans AC, McKinstry RC, Almli CR, 2009) using nonlinear transformation implemented in ANTs v2.1.0 (Avants, Epstein, Grossman, & Gee, 2008). Functional data were motion corrected using MCFLIRT v5.0.9. This was followed by co-registration to the corresponding T1 weighted volume using boundary based registration with nine degrees of freedom, implemented in FreeSurfer v6.0.0. Motion correcting transformations, T1 weighted transformation and MNI template warp were applied in a single step using antsApplyTransformations v2.1.0 with Lanczos interpolation. Three tissue classes were extracted from the T1 weighted images using FSL FAST v5.0.9. Voxels from cerebrospinal fluid and white matter were used to create a mask in turn used to extract physiological noise regressors using aCompCor. Mask was eroded and limited to subcortical regions to limit overlap with grey matter, and six principal components were estimated. Framewise displacements was calculated for each functional run using Nipype implementation. For more details of the pipeline using fMRIprep see http://fmriprep.readthedocs.io/en/1.0.0-rc8/workflows.html.

We created confound files for each scan (each run of each task of each session of each participant), with the following measurements: standard deviation of the root mean squared (RMS) intensity difference from one volume to the next (DVARS), absolute DVARS values, voxelwise standard deviation of DVARS values and six motion parameters (translational and rotation, each in three directions). We added a single time point regressor (a single additional column) for each volume with framewise-displacement value larger than 0.9, in order to model out volumes with extensive motion (i.e. scrubbing). Scans with more than 15% scrubbed volumes were excluded from analysis, resulting in one excluded participant from the analysis of the first session’s probe task. The confounds.tsv files (FSL’s format) can be found with the data shared on OpenNeuro.

### fMRI analysis

Imaging analysis was performed using FEAT (fMRI Expert Analysis Tool) v6.00, part of FSL (FMRIB’s Software Library) (Smith et al., 2004).

#### Univariate imaging analysis - passive viewing

The functional data from the passive viewing task were used to examine the functional changes underlying the behavioral change of preferences following CAT in the short- and long-term. We used a general linear model (GLM) with 13 regressors: Eight regressors modelling each item type (high-value Go items, high-value NoGo items that were included in the probe task, high-value NoGo items that were not included in the probe task, high-value ‘sanity check’ items, and the same four regressors for low-value items); the four regressors which are relevant to the Go / NoGo probe comparisons (high-value Go items, low-value Go items, high-value NoGo items that were included in the probe task and low-value NoGo items that were included in the probe task), with the same onsets and duration, and a parametric modulation by the mean-centered proportion of trials each item was chosen in the subsequent probe phase (the number of trials each item was chosen during the subsequent probe divided by the number of probe trials including this item, mean-centered) and one regressor for all items with a parametric modulation by the mean-centered WTP values acquired from the first BDM auction (which was added to control for initial WTP differences). These 13 regressors were convolved with the canonical double-gamma hemodynamic response function, and their temporal derivatives were added to the model. We further included at least nine motion regressors as confounds, as described above. We estimated a model with the above described GLM regressors for each passive viewing run of each participant in a first level analysis.

In the second level analysis (fixed effects), runs from the same session were averaged and compared to the other session. Two second level contrasts were analyzed separately: after compared to before CAT and follow-up compared to before CAT.

All second level analyses of all participants from after minus before or from follow-up minus before CAT were then inputted to a group level analysis (mixed effects), which included two contrasts of interest: One with the main effect (indicating group mean) and one with the mean centered probe effect of each participant (the demeaned proportion of choosing Go over NoGo items during the subsequent probe for the relevant pair type, i.e. either high-value, low-value or all probe pairs). The second contrast was used to test the correlation between the fMRI activations and the behavioral effect across participants (correlation with the behavioral effect across participants). The design.fsf files (FSL’s format) for each participant, session, task and analysis level can be found with the data shared on OpenNeuro.

All reported group level statistical maps were thresholded at Z > 2.3 and cluster-based Gaussian Random Field corrected for multiple comparisons at the whole-brain level with a (corrected) cluster significance threshold of *P* = 0.05 (Worsley, 2001).

Since we found a stronger behavioral effect for high-value items, similarly to previous cue-approach samples with snack food items (Salomon et al., 2018; Schonberg et al., 2014), we focused our analyses on the contrasts for high-value items: high-value Go items, high-value Go items modulated by choice and high-value Go minus high-value NoGo items. For completeness, we report the results of these contrasts with low-value items (low-value Go items, low-value Go items modulated by choice and low-value Go minus low-value NoGo items), as well as a direct comparison between high-value and low-value items (i.e. high-value Go minus low-value Go items).

#### Univariate imaging analysis – probe

Imaging analysis of the probe data was similar to previous imaging studies with CAT (Bakkour et al., 2017; Schonberg et al., 2014). We included 16 regressors in the model (in addition to at least nine motion regressors as described above), based on the initial value of the probe pair (high / low) and the choice outcome (participant chose the Go / NoGo item), resulting in four regressors (high-value chose Go / high-value chose NoGo / low-value chose Go / low-value chose NoGo) without parametric modulation; the same four regressors with a parametric modulation across items by the mean-centered proportion of choices of the specific item during the entire probe phase; the same four regressors with a parametric modulation by the WTP difference between the two presented items; one regressor for all “sanity-check” trials; one regressor for all missed trials; and two regressors accounting for response time differences (one regressor with a modulation of the demeaned response time across trials for each value category).

Since our behavioral effect was stronger for high-value items (similar to previous cue-approach samples with snack food items), we focused our analysis on the contrasts for high-value chose Go, high-value chose Go modulated by choice and high-value chose Go minus high-value chose NoGo. Similar to analyses of the passive viewing task, we estimated a first level GLM for each run of each participant. We then averaged the four runs of each probe (after / follow-up) of each participant in a second-level analysis. Finally, we ran a group level analysis as described above, with one contrast for the mean group effect and one contrast for the demeaned probe effect across participants (correlation with the behavioral effect across participants).

Some of the probe runs were excluded from the imaging analysis (i.e. not included in the second level analysis of the specific participant) because one of the regressors was empty or because the parametric modulator of Go item choices was zeroed out, resulting in a rank-deficient design matrix. This happened, for example, when a participant chose high-value Go over high-value NoGo items on all trials of a specific run. Participants who did not have at least one full valid block (out of two probe blocks, each probe including one presentation of each probe pair) without any empty regressors or zeroed modulators for Go items, were excluded from the probe imaging analysis. In order to minimize the number of excluded runs and participants, we did not exclude runs or participants due to a zeroed modulator of NoGo items choices, but rather decided not to use the contrasts including modulation by choice of trials where NoGo items were chosen. Overall, one participant was excluded from the imaging analysis of the probe from both the after and follow-up sessions and two more were excluded each from one of the sessions, based on regressors causing rank-deficient matrices (in addition to the one participant that was excluded from the first session due to excessive motion, as described above). Thus, a total of 33 (out of 36) participants were included in the imaging analysis of the probe after training (out of which for 28 participants no run was excluded, for four participants one run was excluded and for one participants two runs-one block-were excluded), and 25 (out of 27) participants were included in the imaging analysis of the follow-up probe (out of which for 21 participants no run was excluded and for four participants one run was excluded).

#### Small volume correction (SVC) analysis

We pre-hypothesized (and pre-registered) that value, attention and memory-related brain regions will be associated with the behavioral change following CAT: Prefrontal cortex, dorsal parietal cortex and medial-temporal lobe, respectively (https://osf.io/q8yct/). Thus, in addition to the whole-brain analyses described above for the passive viewing and probe tasks, we ran similar group level analyses once for each of these pre-hypothesized regions (bilateral hippocampus, bilateral SPL and vmPFC), with a mask containing the voxels which were part of the region. All masks were based on the Harvard-Oxford atlas (see **Supplementary Figure 2**), anatomical regions for the vmPFC mask were based on those used in previous CAT studies (Bakkour et al., 2017; Schonberg et al., 2014). The masks are shared with the data on OpenNeuro.

## Results

### Behavioral probe results

#### After CAT

As expected from previous studies (Bakkour et al., 2016, 2017; Salomon et al., 2018; Schonberg et al., 2014), participants significantly preferred Go over NoGo items in high-value probe choices (mean = 0.590, SE = 0.032, Z = 2.823, *P =* 0.002, one-sided logistic regression) and marginally also in low-value probe choices (mean = 0.561, SE = 0.038, Z = 1.639, *P =* 0.051; Figure 2). The proportion of Go items choices was significantly higher for high-value compared to low-value items (indicating a differential effect of CAT on preference for stimuli of the two value categories; Z = 2.184, *P* = 0.015, one-sided logistic regression).

**Figure 2.**
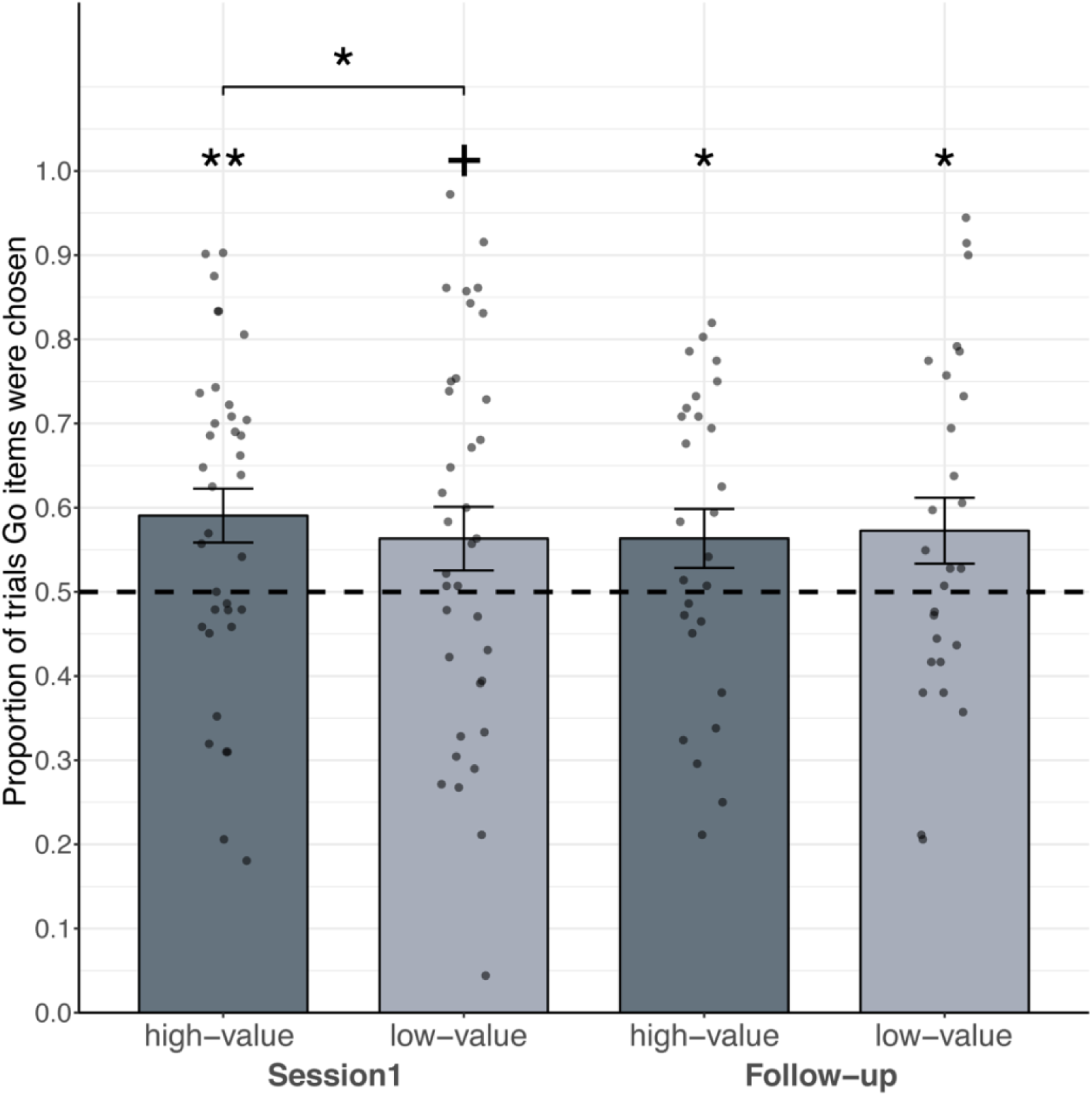
Behavioral results of Go choices during probe: *Mean proportion of trials in which participants chose Go over NoGo items are presented for high-value (dark gray) and low-value (light gray) probe pairs, for each session. Means of the single participants are presented with dots over each bar. The dashed line indicates chance level of 50%, error bars represent standard error of the mean. Asterisks reflect statistical significance in a one-tailed logistic regression analysis. Asterisks above each line represent proportions higher than chance (log-odds = 0, odds-ratio = 1). Asterisks above pairs of bars represent differential effect between the two value categories; +P < 0.1, *P < 0.05, **P < 0.005.*

#### One-month follow-up

One month following training (mean = 30.26 days, SD = 9.93 days, N = 27) participants significantly chose Go over NoGo items in both high-value (mean = 0.563, SE = 0.035, Z = 1.854, *P =* 0.032) and low-value (mean = 0.572, SE = 0.039, Z = 1.948, *P =* 0.026) probe trials. There was no differential effect between high and low-value items in this session (Z = 0.622, *P* = 0.267).

### Behavioral recognition results

There was a prominent ceiling effect in participants’ performance on the recognition task, in both sessions. This ceiling effect did not allow us to reveal differences in memory for Go compared to NoGo items. For full description and statistics see Supplementary analysis: behavioral recognition results.

#### Immediately after CAT

The mean hit rate across participants was 99.29% (SD = 2.08%), mean correct rejection rate of 94.16% (SD = 5.22%) and mean d’ = 3.921 (SD = 0.547). We did not find significant differences in hit rate or in RT between Go and NoGo items.

#### One-month follow-up

It should be noted again that the recognition task in the follow-up session was performed immediately after the passive viewing and probe tasks. Therefore, results reflect within-session memory, and not long-term memory effects from the first session. The mean hit rate was 97.81% (SD = 2.37%), mean correct rejection rate 94.26% (SD = 7.77%) and mean d’ was 3.79 (SD = 0.715). Again, we did not find significant differences in hit rate or in RT between Go and NoGo items.

### Behavioral auction results

Similar to previous results regarding the change in WTP following CAT (Schonberg et al., 2014), we observed a general trend of regression to the mean - i.e. while WTP for high-value items decreased, WTP for low-value items increased. This trend was descriptively weaker for Go compared to NoGo items in both sessions (immediately after compared to before CAT and one-month after compared to before CAT) and both value-level (high-value and low-value items), but these differences were very small and were not significant. For full description and statistics see Supplementary analysis: behavioral auction results.

### Imaging results

Behavioral results with snack food items from previous studies (Bakkour et al., 2017; Salomon et al., 2018; Schonberg et al., 2014) and from the current study demonstrated a differential pattern of the change of preferences across value levels: Preference modifications were more robust for high-value compared to low-value items; therefore, we chose to focus on the functional changes in the representation of high-value items. Functional changes in the representation of low-value items, as well as the direct comparison between the change in high-value and low-value items, are also reported and are presented in the supplementary material.

We further tested two kinds of relations between the behavioral effect and the neural response: *Modulation across items*, meaning that the change in activity was stronger for items that were later more preferred during the subsequent probe phase (within-participant first-level parametric modulation); and *correlation across participants*, meaning that the change in activity was stronger for participants that later showed a stronger behavioral probe effect, quantified as a higher ratio of choosing Go over NoGo items (between-participants group-level correlation). Finally, for a subset of three pre-hypothesized and pre-registered regions (vmPFC, hippocampus and superior parietal lobule) we performed a small volume correction (SVC) analysis (see methods and materials).

All reported group level statistical maps were thresholded at Z > 2.3 and cluster-based Gaussian Random Field corrected for multiple comparisons at the whole-brain level with a (corrected) cluster significance threshold of *P* = 0.05 (Worsley, 2001). Unthresholded and thresholded images of all contrasts presented here are shared on NeuroVault (Gorgolewski et al., 2015) (https://neurovault.org/collections/TTZTGQNU/)

### Passive viewing imaging results

To investigate the functional changes in the response to individual items following CAT, we scanned participants with fMRI while they were passively viewing the items. Participants completed this task before, after and one month following CAT (N = 36 before and immediately after, and N = 27 after one month).

#### Immediately after versus before CAT (Figure 3, for description of all activations see Supplementary Table 2)

BOLD activity while passively viewing high-value Go compared to passively viewing high-value NoGo items was increased after compared to before CAT in the left and right occipital and temporal lobes (Figure 3a), along the ventral visual processing pathway (Goodale & Milner, 1992). Results of the SVC analyses revealed enhanced BOLD activity during passive viewing of high-value Go items after compared to before CAT in the vmPFC (Figure 3b).

**Figure 3.**
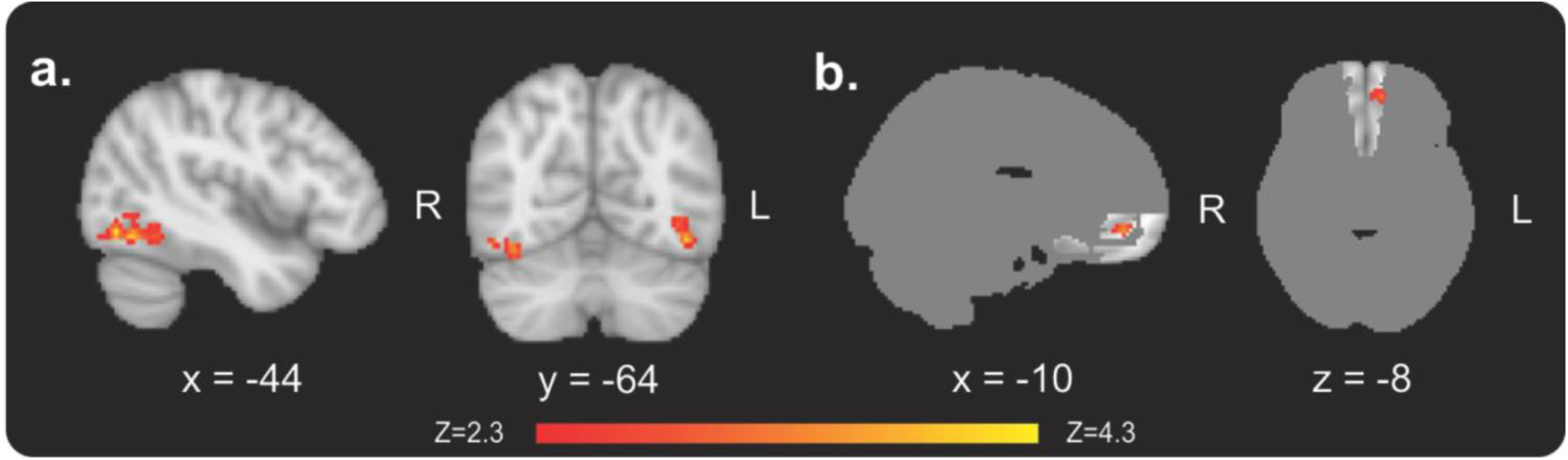
fMRI results from the passive viewing task after compared to before CAT: ***(a)*** *Enhanced BOLD activity in bilateral occipito-temporal regions, for high-value Go compared to high-value NoGo items (whole-brain analysis).* ***(b)*** *Enhanced BOLD activity in the vmPFC in response to high-value Go items (Small volume corrected results; the mask used to correct for multiple comparisons is presented on a dark grey brain silhouette). For description of all activations see Supplementary Table 2.*

Bold activity while passively viewing low-value Go compared to passively viewing low-value NoGo items was stronger immediately after compared to before CAT in the temporo-occipital part of the left middle temporal gyrus, left superior lateral occipital cortex, left posterior supramarginal / angular gyrus, middle PFC and cerebellum (see Supplementary Figure 3). The significant cluster for the low-value items in the temporo-occipital visual cortex was smaller, as well as more anterior and lateral, compared to the significant cluster for the high-value items. SVC analysis in the vmPFC for low-value items immediately after compared to before CAT did not reveal any significant results.

A direct comparison between the changes for high-value Go compared to low-value Go items, as well as a comparison between the difference of high-value Go minus high-value NoGo items and the difference of low-value Go minus low-value NoGo items, revealed no significant clusters.

#### One-month follow-up versus before (Figure 4, for description of all activations see Supplementary Table 3)

BOLD activity in the vmPFC was enhanced one month after compared to before CAT in a SVC analyses (Figure 4a), similarly to the short-term change. In addition, BOLD activity in the left orbitofrontal cortex (OFC) in response to high-value Go items was positively modulated by the choice effect across items in the follow-up compared to before CAT (whole-brain analysis; Figure 4b). SVC analyses revealed that BOLD activity in response to high-value Go items in the right anterior hippocampus was positively modulated by the choice effect across items in the follow-up compared to before training (Figure 4c), while BOLD activity in response to high-value Go minus high-value NoGo items in the right SPL was negatively correlated with the choice effect across participants in the follow-up compared to before training (Figure 4d).

**Figure 4.**
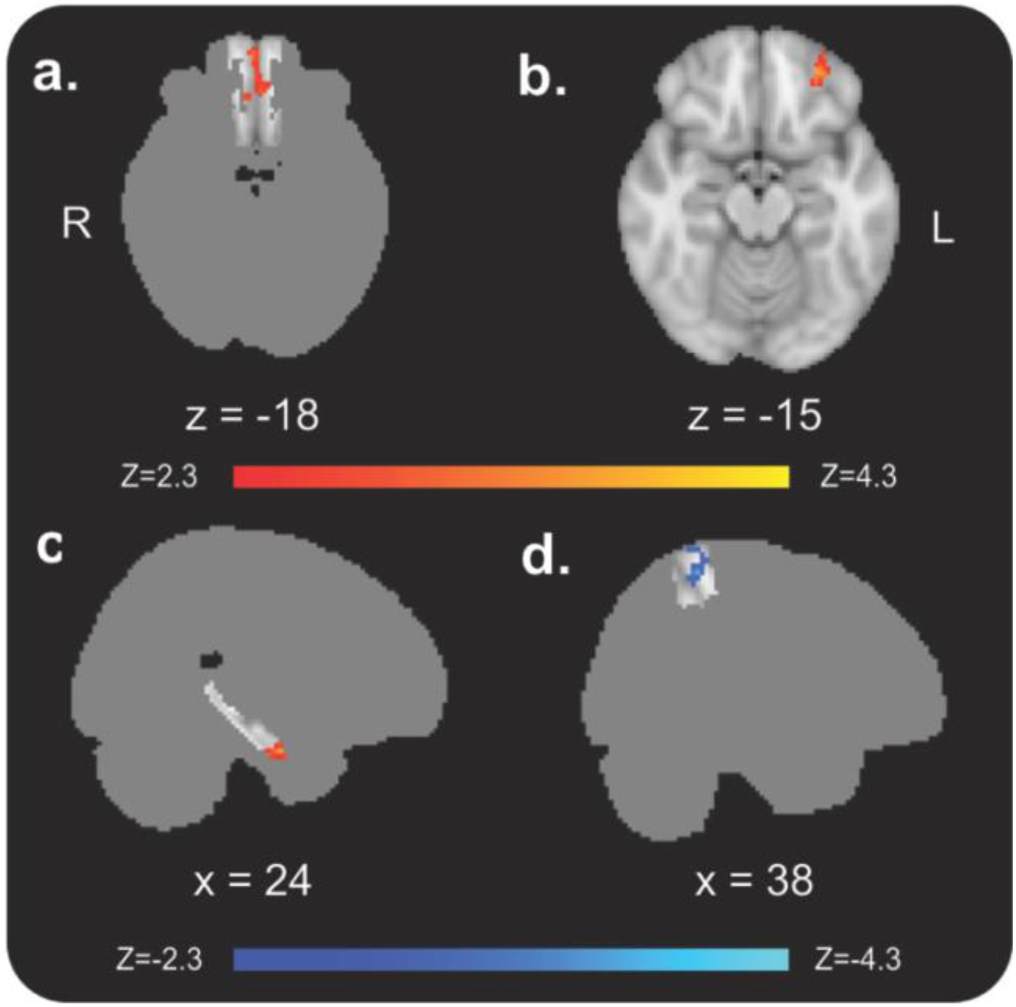
fMRI results from the passive viewing task in the one-month follow-up compared to before CAT: ***(a)*** *Enhanced BOLD activity in response to high-value Go items in the vmPFC (small volume corrected).* ***(b)*** *BOLD activity in response to high-value Go items in the left OFC was positively modulated by the choice effect across items (whole-brain analysis).* ***(c)*** *BOLD activity in response to high-value Go items in the right anterior hippocampus was positively modulated by the choice effect across items (small volume corrected).* ***(d)*** *BOLD activity in response to high-value Go minus high-value NoGo items in the right SPL was negatively correlated with the choice effect across participants (small volume corrected). The masks used to correct for multiple comparisons in the small volume correction (SVC) analyses are presented on a dark grey brain silhouette. For description of all activations see Supplementary Table 3.*

When observing the same contrasts for low-value items, we did not find significant changes in BOLD activity during passive viewing one month following compared to before CAT. A direct comparison between changes for high-value Go compared to low-value Go items also did not reveal significant differences.

### Probe imaging results

To investigate the functional response during choices, we scanned participants with fMRI while they completed the probe (binary choices) phase, as was done in previous studies (Bakkour et al., 2017; Schonberg et al., 2014). Participants completed the probe task immediately after CAT (N = 33). In the current study, we also scanned for the first time the probe session in the one-month follow-up (N = 25).

#### Immediate Probe (Figure 5a-e, for description of all activations see Supplementary Table 4)

BOLD activity was stronger during choices of high-value Go over high-value NoGo items compared to choices of high-value NoGo over high-value Go items in bilateral visual regions and bilateral central opercular cortex and Heschl’s gyrus (Figure 5a). In addition, BOLD activity in the striatum while choosing high-value Go compared to choosing high-value NoGo items after CAT was negatively correlated with the choice effect across participants (the ratio of choosing high-value Go items during probe; Figure 5b) and negatively modulated by the choice affect across items (Figure 5c). SVC analysis revealed that BOLD activity in the right SPL while choosing high-value Go items after CAT was negatively correlated with the choice effect across participants (Figure 5d) and negatively modulated by the choice effect across items (Figure 5e).

**Figure 5.**
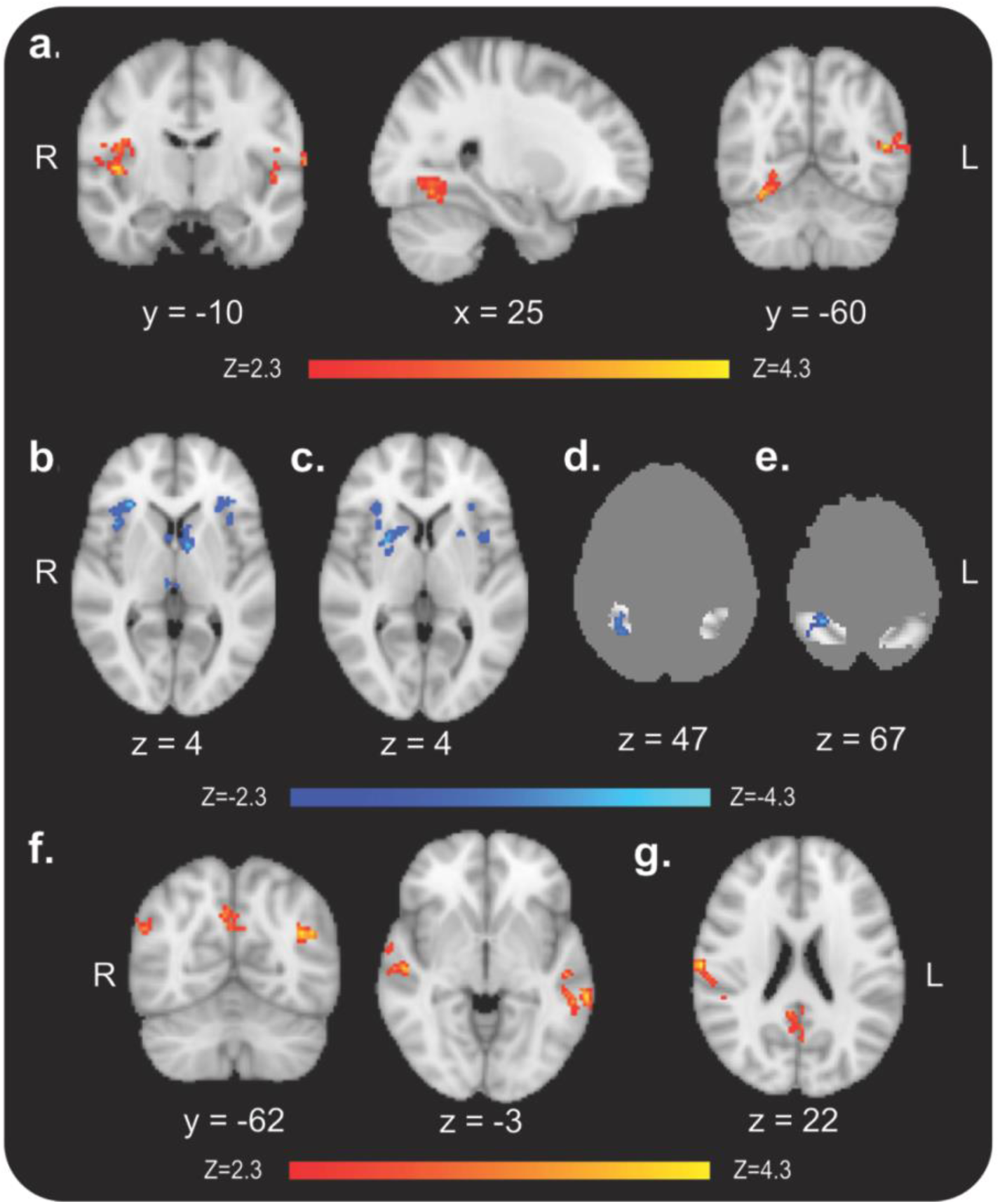
fMRI results from the probe task, immediately after and 30 days following CAT: ***(a)*** *Enhanced BOLD activity during choices of high-value Go compared to choices of high-value NoGo items after CAT in bilateral visual regions and bilateral central opercular cortex and Heschl’s gyrus (whole-brain analysis).* ***(b)*** *BOLD response after CAT was negatively correlated with the choice effect across participants and* ***(c)*** *negatively modulated by the choice effect across items, during choices of high-value Go over high-value NoGo items in the striatum as well as other regions (whole-brain analysis).* ***(d)*** *BOLD response after CAT was negatively correlated with the choice effect across participants and* ***(e)*** *negatively modulated by the choice effect across items, during choices of high-value Go compared to high-value NoGo items, in the right SPL (small volume corrected).* ***(f)*** *Bold activity during choices of high-value Go items (whole-brain analyses) was positively modulated by the choice effect across items in the precuneus, bilateral superior occipital cortex and bilateral middle and superior temporal gyrus and* ***(g)*** *positively correlated with the choice effect across participants in the precuneus/posterior cingulate cortex (PCC) and right post-central gyrus. The masks used to correct for multiple comparisons in the small volume correction (SVC) analyses are presented on a dark grey brain silhouette. For description of all activations see Supplementary Table 4 and Supplementary Table 5.*

There were no regions with significant stronger BOLD activity during choices of low-value Go items compared to choices of low-value NoGo items after CAT. In the superior division of the left lateral occipital cortex, the difference between activity during choices of low-value Go items and activity during choices of low-value NoGo items was negatively correlated with the choice effect across participants (see Supplementary Figure 4a). In addition, BOLD activity in the left and right Heschl’s gyrus / central opercular cortex and in the left precuneous, was stronger immediately after CAT during choices of high-value Go items compared to choices of low-value Go items (see Supplementary Figure 4b).

#### One-month follow-up Probe (Figure 5f-g, for description of all activations see Supplementary Table 5)

BOLD activity in the precuneus, bilateral superior occipital cortex and bilateral middle and superior temporal gyrus while choosing high-value Go items in the follow-up probe was positively modulated by the choice effect across items (Figure 5f). BOLD activity in the precuneus/posterior cingulate cortex (PCC) and right post-central gyrus while choosing high-value Go items in the follow-up probe was positively correlated with the choice effect across participants (Figure 5g). There were no significant results for these contrasts for low-value items. However, BOLD activity in the left and right occipital poles was stronger one month after CAT during choices of high-value Go items compared to choices of low-value Go items (see Supplementary Figure 4c).

## Discussion

In the current work, we set out to examine the neural mechanisms underlying non-reinforced behavioral change following CAT, in the short- and long-term. We introduced a novel passive viewing task to study the functional plasticity of response to single items before, after and one month following CAT. Prior to data analysis, we predicted and pre-registered that the underlying neural mechanisms will involve memory, attention and value-related brain regions.

The behavioral results obtained in the current study (see Figure 2) replicated previous results, demonstrating enhanced preferences towards high-value cued (high-value Go) compared to high-value non-cued (high-value NoGo) items following CAT (Bakkour et al., 2016, 2018, 2017; Salomon et al., 2018; Schonberg et al., 2014; Veling et al., 2017; Zoltak et al., 2017). Similar to previous studies (Schonberg et al., 2014), participants completed a BDM auction task at the beginning of the experiment as well as at the end of each session. In the current study, we found no differences in the regression to the mean for both Go and NoGo items. Previous CAT studied identified regression to the mean for both Go and NoGo items (Schonberg et al., 2014). One previous study found regression to the mean to be weaker for Go compared to NoGo items, but a different sample in the same study did not replicate this effect (Schonberg et al., 2014). These findings suggest that testing WTP difference following CAT is not sensitive enough to behaviorally reveal the value change, while a forced choice paradigm between similarly valued stimuli can detect value modification effects more reliably.

The fMRI results of this study suggest the involvement of several neural components in preference modification induced by CAT.

### Bottom-up mechanisms in the short-term

Examining the neural response for high-value Go compared to high-value NoGo items following CAT revealed enhanced processing in ventral visual regions (Figure 3a). We also found an indication for enhancement of visual processing in response to low-value Go items following CAT (Supplementary Figure 3). This enhancement was in a smaller region, which might be related to the weaker behavioral effect for low-value items. There were no significant results in a direct comparison between the change of response to high-value Go versus low-value Go items. However, it should be noted that our study was not well powered for this comparison, since we focused our analyses on the high-value items, for which there was a stronger behavioral change, both in this study and previous ones with snack food items (Salomon et al., 2018; Schonberg et al., 2014). Eye-gaze recoded during the passive viewing task from a subset of participants did not reveal longer gaze duration on paired items (see Supplementary analysis: eye-tracking) immediately after training and thus is probably not the reason for the enhanced fMRI signal for high-value Go over NoGo.

We show here for the first time that activity in high-level visual processing occipito-temporal cortex is related to subjective values, without external reinforcements. Activity in low and high-level visual regions was previously shown to be related to past rewards (Serences, 2008), but not to subjective values in the absence of reinforcements as in the current study. We suggest that the functional changes in visual regions reflect modifications in the bottom-up perceptual representation of the paired items (Grill-Spector, 2003; Ishai, Ungerleider, Martin, & Haxby, 2000). In the short-term, the enhanced bottom-up processing and representation change of individual Go items lead to enhanced value-related processing and enhanced preferences towards these items during choices.

### Long-term maintenance via memory processes

In the one-month follow-up, hippocampal activity during passive viewing was stronger for high-value Go items that were later chosen more during the subsequent probe (see Figure 4c). This finding suggests, as predicted and pre-registered, that memory-related processes supported the long-term maintenance of the behavioral effect. Importantly, it is the first demonstration, to the best of our knowledge, of the relation between value-based decision-making and memory during passive viewing of items and not during a memory or a choice task. The immediate bottom-up perceptual processing enhancement putatively affected the encoding and accessibility of paired items and their related associations in memory for the long-term. Thus, in the choice phase, more accessible positive associations for the paired items accumulate faster to choices (Shadlen & Shohamy, 2016).

Although participants completed a behavioral recognition task at the end of the experiment, this experiment was not designed to test behavioral memory modifications. Therefore, the recognition task was performed immediately after the rest of the tasks. This design resulted in a ceiling effect of over 99% hit rate across participants, masking any memory differences between Go and NoGo items. Differences in RT were also not optimal for testing, as the task was self-paced. Moreover, in the follow-up session, the recognition task was again performed following the probe task, therefore long-term memory was not actually tested. Thus, our behavioral recognition data do not allow us to behaviorally test for memory involvement in the short- or long-term effect of CAT. Further behavioral studies, designed specifically to test for memory modifications, are required to explore the involvement of memory processes in the behavioral change following CAT, both in the short- and long-term.

### Decreased top-down attention

Participants with overall greater long-term behavioral change demonstrated reduced change of response to high-value Go items in the follow-up compared to before CAT in the right SPL (i.e. activity was negatively correlated with the choice effect across participants in the long-term; see Figure 4d). This finding suggests, as pre-registered, reduced involvement of top-down parietal mechanisms during passive viewing of Go compared to NoGo items (Culham & Kanwisher, 2001). Decreased top-down attentional mechanisms may underlie the impulsive-like nature of the preference bias towards Go items (Veling et al., 2017).

### Enhanced value response, putatively related to enhanced bottom-up over top-down processing

Value change is reflected in enhanced neural response of the vmPFC to high-value Go items, both immediately after CAT (see Figure 3b) and in the one-month follow-up (see Figure 4a). In the one-month follow-up, value change was further reflected in the OFC, where activity was stronger while passively viewing high-value Go items that were later chosen more during the subsequent probe phase (see Figure 4b). These findings may indicate, as predicted and pre-registered, a long-lasting value change signature of individual items not during choices (Lebreton, Jorge, Michel, Thirion, & Pessiglione, 2009; Levy, Lazzaro, Rutledge, & Glimcher, 2011). Overall, these results reveal for the first time an item-level value change during passive viewing (Levy et al., 2011; Serences, 2008), in line with previous findings of enhanced activity in the vmPFC during binary choices of more preferred high-value Go items (Schonberg et al., 2014). In addition, this is the first time, to the best of our knowledge, that such enhancement in value-related prefrontal regions was found one month following a behavioral change paradigm.

We suggest that the joint effect of the three above described components provides a shift towards enhanced bottom-up over top-down processes, resulting in non-externally reinforced behavioral change which can be maintained for a long period of time. Based on the findings of the current study, we propose that the low-level association of visual, auditory and motor systems during training modifies valuation of items via a network including: enhanced bottom-up perceptual processes in the immediate short-term, which is translated to long term maintenance by memory enhancement and decrease of top-down attentional control. This leads to long-lasting behavioral change. In Figure 6 we provide a putative outline of the suggested dynamics of this preference change process. Although the proposed model dynamics are based on reverse inference from the current work data, its principle components (i.e. - memory, top-down attention and value-related mechanisms) were predicted and pre-registered prior to analyses. The methods and results of our current study do not lend themselves to directly testing this suggested mechanism using current available methods. Future studies should test the validity and reproducibility of these results using independent data.

**Figure 6.**
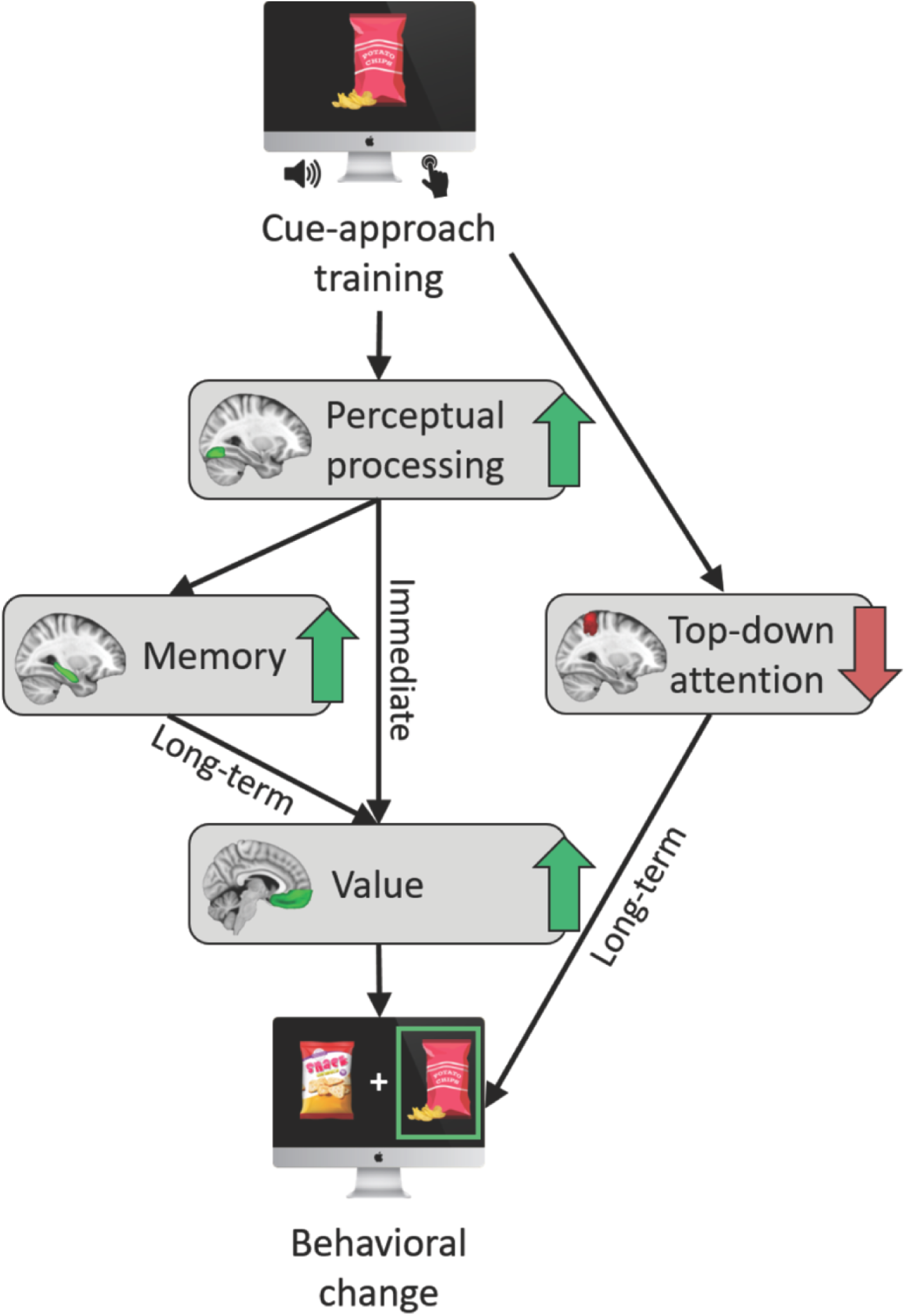
Suggested dynamics of the preference change. *We propose that training leads to enhanced perceptual processing, which leads to value enhancement in the short-term and thus to immediate behavioral change. The enhanced perceptual processing putatively enhances memory activation and accessibility, which drives the long-lasting behavioral change. In addition, the involvement of top-down attention is reduced following training, further enhancing the long-term behavioral change.*

### Functional activity during binary choices also reflect enhanced bottom-up and decreased top-down mechanisms of preferences modification

Functional MRI responses during binary choice also resonate the proposed dynamics for the non-reinforced behavioral change dynamics we are proposing (Figure 6), demonstrating enhanced perceptual processing in the short-term and involvement of memory processes in the long-term, as well as decreased top-down attention mechanisms.

When participants chose high-value Go over high-value NoGo items, activity in perceptual regions - both visual and auditory - was enhanced (see Figure 5a). These findings suggest that in the short-term, retrieval of the low-level visual and auditory associations constructed during training, were associated with choices of Go items. Thus, functional response during choices further supports the involvement of enhanced bottom-up processing in the immediate non-reinforced modification of preferences.

In the follow-up session, choices of high-value Go over NoGo items were associated with enhanced BOLD activity in the precuneus and posterior cingulate cortex (PCC; see Figure 5f-g), which have been related to episodic memory retrieval and are also considered to be part of the default mode network (Fransson & Marrelec, 2008; Sestieri, Corbetta, Romani, & Shulman, 2011; Shallice et al., 1994). This provides additional support for the central role of memory processes in the long-term retention of the behavioral change.

Finally, activity in the right SPL during choices of Go items was negatively modulated by the choice effect across items as well as negatively correlated with the choice effect across participants (though in a more posterior and inferior region, see Figure 5d-e), indicating that top-down attentional mechanisms were less involved during choices of Go compared to choices of NoGo items (Culham & Kanwisher, 2001). These findings demonstrate again that reduced top-down mechanisms are involved in the behavioral change following CAT.

We were not able to replicate previous results showing enhanced activity in the vmPFC during choices of Go items that were chosen more overall (Bakkour et al., 2017; Schonberg et al., 2014). These previous results were found for high-value Go items when the group’s behavioral effect of choosing high-value Go items was significant but weak relative to other samples (study 3 in Schonberg et al., 2014). Similar results were found for choices of low-value Go compared to choices of low-value NoGo items, and not for choices of high-value Go items, when the behavioral effect was strong for high-value items and weak for low-value items (Bakkour et al., 2017). Therefore, a possible account for the lack of replication of these findings in the current study is that this contrast of modulation across items depends on the variance of the choice effect across items, which seems to be smaller in the current study compared to previous samples that found this effect.

Another novel finding was that activity in the striatum was negatively correlated with choices of high-value Go over NoGo items (see Figure 5b-c). The striatum is known to be involved in reinforcement learning and habit-based learning (Daniel & Pollmann, 2014; Yin & Knowlton, 2006). These findings potentially suggest that cue-approach training shifted the process of goal-directed decision-making during binary choices to rely more on bottom-up non-reinforced mechanisms. This is the first study with CAT to observe these effects during probe and thus it remains to be replicated in future studies.

Overall, neural activity during binary choices support our suggested proposal of non-reinforced behavioral change (Figure 6), demonstrating similar patterns to these shown in the passive viewing task: enhanced perceptual processing in the short-term, long-term manifestation of the behavioral change through memory-related mechanisms, and reduced top-down involvement (here both in the short and long-term).

## Conclusions

Research of value-based decision-making and behavioral change interventions focused on top-down mechanisms such as self-control or external reinforcements as the main means to change preferences (Rangel, Camerer, & Montague, 2008; Wood & Neal, 2016). The cue-approach training paradigm has been shown to change preferences using the mere association of images of items with a cued speeded button response without external reinforcements. The paradigm is highly replicable with dozens of studies demonstrating the ability to change behavior for months with various stimuli and cues (Bakkour et al., 2016, 2018, 2017; Salomon et al., 2018; Schonberg et al., 2014; Veling et al., 2017; Zoltak et al., 2017).

Current interventions that rely on reinforcement and self-control fail to change behavior for the long-term. Our findings emphasize the importance and great potential of targeting bottom-up rather than top-down mechanisms to induce long-lasting behavioral change. Our results further emphasize the involvement of memory processes in value-based decision-making (even in the absence of choice or memory task) and its relevance to the durability of the behavioral change. We present a suggested model for the dynamics underlying this change. Our current findings can lead to new theories relating perceptual processing, memory and attention to preferences and decision-making. They hold promise for new long-term behavioral change interventions targeting this novel pathway for value change based on bottom-up mechanisms, which can lead to long lasting change and thus improve the quality of life for people around the world.

## Acknowledgments

We thank Dr. Jeanette Mumford for her invaluable statistics advices.

