## Supplementary materials for "Enhanced bottom-up and reduced top-down fMRI activity is related to long-lasting non-reinforced behavioral change"

### Supplementary Material

#### Supplementary analysis: behavioral recognition results

**Immediately after CAT:** As noted in the methods section, we excluded trials with RTs longer than three standard deviations above the mean, for each version of the task. Overall, seven trials were excluded from the results of the binary version (with RT longer than 3.211 secs) and 38 trials were excluded from the results of the confidence version (with RT longer than 5.536 secs). There was a prominent ceiling effect in participants' performance on the recognition task with a mean hit rate of 99.29% (SD = 2.08%) and mean correct rejection rate of 94.16% (SD = 5.22%); mean d' = 3.921 (SD = 0.547). The mean RT was 1.467 (SD = 0.1389) seconds for hits, 1.828 (SD = 1.541) seconds for misses, 1.519 (SD = 0.420) seconds for correct rejections and 2.220 (SD = 0.924) seconds for false alarms.

Hit rate in the old/new recognition task was only descriptively higher for Go (mean = 99.77%, SD = 1.39%) compared to NoGo (mean = 99.05%, SD = 3.37%) items (logistic regression, two-sided $P$ = 0.181, z = 1.339). RTs in the recognition task were also only descriptively shorter for Go (mean = 1.419 secs, SD = 0.40 secs) compared to NoGo (mean = 1.453 secs, SD = 0.41 secs) items (linear regression, two-sided $P$ = 0.404, t = - 0.836).

**One-month follow-up:** It should be noted again that the recognition task in the follow-up session was performed after the passive viewing and probe tasks. Therefore, results reflect within-session memory, and not long-term memory effects from the first session. In the follow-up session, the old/new recognition task was performed with the confidence version (n = 23 participants). Thirty trials were excluded from analysis (with RTs longer than 5.088 seconds). There was a prominent ceiling effect in participants' performance in the recognition task ($M_{(Hit)}$ = 97.81%, $SD_{(Hit)}$ = 2.37%; $M_{(Correct\ Rejection)}$ = 94.26%, $SD_{(Correct\ Rejection)}$ = 7.77%; $M_{(d')}$ = 3.79, $SD_{(d')}$ = 0.715). The mean RT was 1.486 (SD = 0.384) seconds for hits, 1.671 (SD = 1.260) seconds for misses, 1.509 (SD = 0.506) seconds for correct rejections and 2.403 (SD = 1.014) seconds for false alarms.

Hit rate in the old/new recognition task was only descriptively higher for Go (mean = 98.16%, SD = 3.58%) compared to NoGo (mean = 97.83%, SD = 3.74%) items (logistic regression, two-sided *P* = 0.747, z = 0.323). RTs in the recognition task were also only descriptively shorter for Go (mean = 1.450 secs, SD = 0.39 secs) compared to NoGo (mean = 1.461 secs, SD = 0.42 secs) items (linear regression, two-sided *P* = 0.822, t = - 0.225).

**Supplementary analysis: behavioral auction results**

Similar to previous results regarding the change in WTP following CAT (Schonberg et al., 2014), we observed a general trend of regression to the mean - i.e. while WTP for high-value items decreased, WTP for low-value items increased.

**Immediately after compared to before CAT:** The regression to the mean was descriptively weaker for Go compared to NoGo items, but the effect was not statistically significant for high-value items (Go items: mean ΔWTP = -0.542 ILS; NoGo items: mean ΔWTP = -0.554 ILS; *P* = 0.918, two-sided linear regression), nor for low-value items (Go items: mean ΔWTP = 0.608 ILS; NoGo items: mean ΔWTP = 0.695 ILS; *P* = 0.482, two-sided linear regression).

**One-month follow-up compared to before CAT:** Similarly, the regression to the mean was descriptively weaker for Go compared to NoGo items, but this difference was not significant for high-value (Go items: mean ΔWTP = -1.070 ILS; NoGo items: mean ΔWTP = -1.124 ILS; *P* = 0.688, two-sided linear regression) nor for low-value items (Go items: mean ΔWTP = 0.737 ILS; NoGo items: mean ΔWTP = 0.843 ILS; *P* = 0.453, two-sided linear regression).

**Supplementary analysis: eye-tracking**

Eye-tracking data were recorded from a subset of participants, using an EyeLink 1000 Plus SR-Research eye-tracker. For the passive viewing task, we had useable eye-gaze data from 10 participants after CAT and 10 participants in the one-month follow-up (we did not obtain eye-tracking data prior to training). Eye-tracking data from the passive viewing task were used to test whether the duration of time spent observing Go items was different from the duration observing NoGo items after CAT. We averaged the time spent looking on Go items and the time spent looking on NoGo items for each participant and performed a paired t-test to test for differences in observation time between Go and NoGo items.

We found no differences in eye-gaze duration between high-value Go and NoGo items during the task, neither after CAT (mean percent of viewing time: high-value Go 75.3%, high-value NoGo 77.4%; *P* = 0.512) nor in the follow-up session (mean percent of viewing time: high-value Go 77.9%, high-value NoGo 74.2%; *P* = 0.266).

**Supplementary figures**

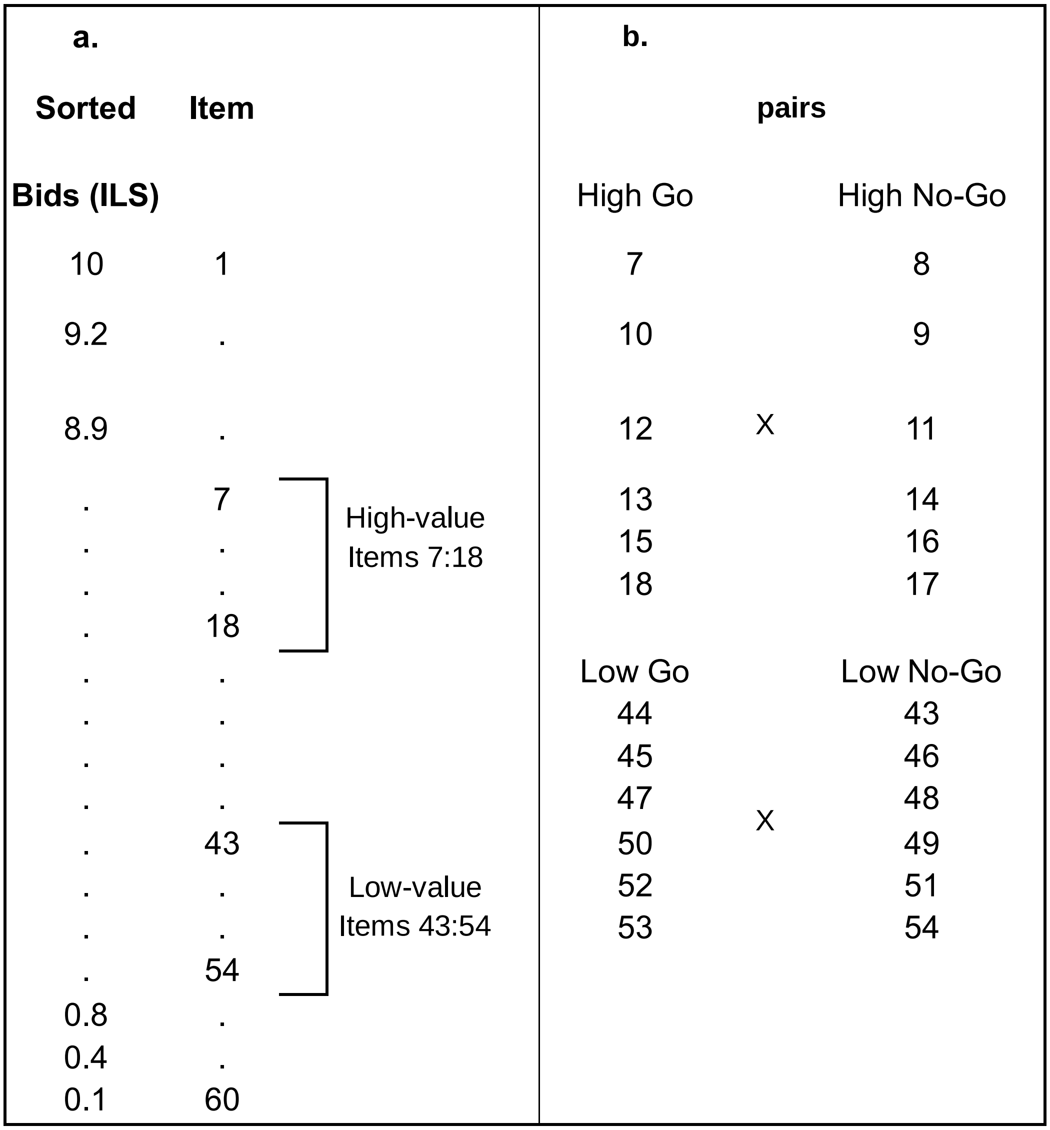


***Supplementary Figure 1. Diagram of the item selection procedure used in this study.***

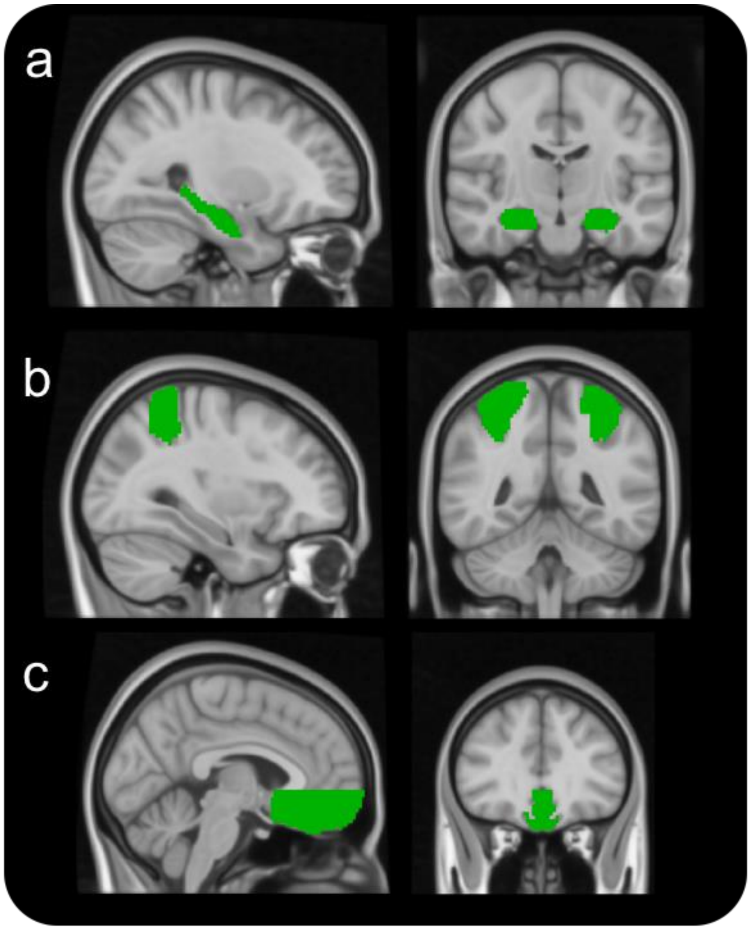


***Supplementary Figure 2. Masks for small volume correction:* *(a)*** *Left and right hippocampus;* ***(b)*** *left and right SPL;* ***(c)*** *vmPFC.*

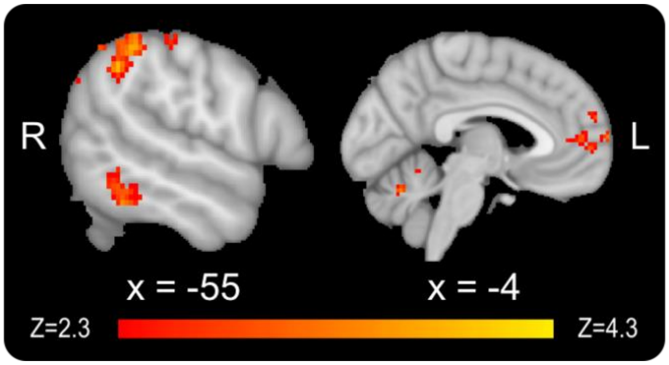


***Supplementary Figure 3. fMRI results from the passive viewing task after compared to before CAT, low-value items:*** *Enhanced BOLD activity in the temporo-occipital part of the left middle temporal gyrus, left superior lateral occipital cortex, left posterior supramarginal / angular gyrus, middle PFC*

*and cerebellum, while passively observing low-value Go compared to low-value NoGo items (whole-brain analysis).*

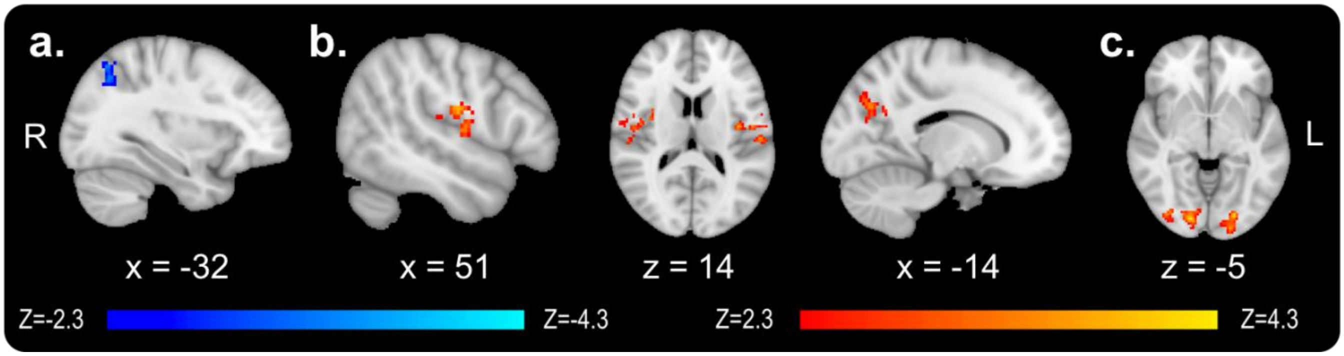


***Supplementary Figure 4. fMRI results from the probe task, immediately after and 30 days following CAT, for low-value items: (a)*** *BOLD activity after CAT was negatively correlated with the choice effect across participants during choices of low-value Go over low-value NoGo items in the left superior lateral occipital cortex (whole-brain analysis).* ***(b)*** *BOLD activity after CAT was stronger during choices of high-value Go items compared to choices of low-value Go items in the left and right Heschl's gyrus / central opercular cortex, as well as in the left precuneous.* ***(c)*** *BOLD activity 30 days after CAT was stronger during choices of high-value Go items compared to choices of low-value Go items in the left and right occipital poles.*

### Supporting tables

| | Brand | Description | | Brand | Description |
|---|---|---|---|---|---|
| **1** | Apropo | Salty snack | **31** | GumiWine | Gummy candy |
| **2** | BabyDoll | Candy | **32** | Halva | Halva |
| **3** | BagaleShtuhim | Salty snack | **33** | HappyHippo | Chocolate |
| **4** | Bamba | Salty snack | **34** | HispusimShatiah | Gummy candy |
| **5** | BambaNugat | Sweet and salty snack | **35** | Hit | Cookies |
| **6** | BambaSweet | Sweet and salty snack | **36** | KashitSour | Gummy candy |
| **7** | BisliBbq | Salty snack | **37** | Keifli | Salty snack |
| **8** | BisliGrill | Salty snack | **38** | KifkefMaklot | Chocolate |
| **9** | BisliOnion | Salty snack | **39** | KinderBuenoBrown | Chocolate |
| **10** | BisliPizza | Salty snack | **40** | KinderBuenoWhite | Chocolate |
| **11** | Bounty | Chocolate | **41** | KinderJoy | Chocolate |
| **12** | Cheetos | Salty snack | **42** | Kitkat | Chocolate |
| **13** | ChocolateParaCookies | Chocolate | **43** | Loacker | Wafer |
| **14** | ChocolateParaMarir | Chocolate | **44** | Mars | Chocolate |
| **15** | ChocolateParaMilk | Chocolate | **45** | Mekupelet | Chocolate |
| **16** | ChocolateParaSucariot | Chocolate | **46** | Mentos | Candy |
| **17** | ClickBalls | Chocolate | **47** | PesekZman | Chocolate |
| **18** | ClickBisquit | Chocolate | **48** | Popco | Sweet and salty snack |
| **19** | ClickBlackWhite | Chocolate | **49** | Shugi | Energy bar |
| **20** | ClickXLbrown | Chocolate | **50** | SkittlesFruits | Candy |
| **21** | ClickXLwhite | Chocolate | **51** | SkittlesSour | Candy |
| **22** | CrunchBisquitBrown | Chocolate | **52** | Smarties | Candy |
| **23** | CrunchBisquitWhite | Chocolate | **53** | Snickers | Chocolate |
| **24** | CrunchShokoVanil | Chocolate | **54** | Taami | Chocolate |
| **25** | DoritosGrill | Salty snack | **55** | Tapuchips | Salty snack |
| **26** | DoritosNatural | Salty snack | **56** | Tictac | Candy |
| **27** | DoritosSourSpicy | Salty snack | **57** | Tortit | Chocolate |
| **28** | Dubonim | Salty snack | **58** | Twist | Wafer |
| **29** | Egozi | Chocolate | **59** | Twix | Wafer |
| **30** | GumiSnakes | Gummy candy | **60** | WerthersOriginal | Candy |

***Supplementary Table 1. Snack food stimuli used in this study.***

*The complete images data set is available online:*

*http://schonberglab.tau.ac.il/resources/snack-food-image-database/*

| Contrast | Cluster | Region | Number of voxels in region | Cluster size | X | Y | Z | Peak z stat |
|---|---|---|---|---|---|---|---|---|
| High-value Go minus NoGo (Fig 3a) | 1 | L Lateral Occipital Cortex, inferior division | 104 | 171 | -44 | -72 | -11 | 4 |
| | | L Inferior Temporal Gyrus, temporooccipital part | 38 | | | | | |
| | | L Temporal Occipital Fusiform Cortex | 19 | | | | | |
| | 2 | R Temporal Occipital Fusiform Cortex | 107 | 192 | 30 | -46 | -8 | 3.82 |
| | | R Lingual Gyrus | 33 | | | | | |
| | | R Occipital Fusiform Gyrus | 22 | | | | | |
| | | R Lateral Occipital Cortex, inferior division | 20 | | | | | |
| High-value Go, small volume correction (vmPFC; Fig 3b) | 1 | L Frontal Medial Cortex | 50 | 98 | -10 | 48 | -11 | 3.44 |
| | | R Frontal Medial Cortex | 33 | | | | | |

***Supplementary Table 2*. *Passive viewing task, after minus before CAT:*** *Regions with significant activation for the imaging contrasts of* ***Figure 3****. For each cluster, the list presents all regions from the Harvard-Oxford atlas that contained at least 10 active voxels within the cluster, as well as the X/Y/Z location for the peak activation in MNI space.*

| Contrast | Cluster | Region | Number of voxels in region | Cluster size | X | Y | Z | Peak z stat |
|---|---|---|---|---|---|---|---|---|
| High-value Go, small volume correction (vmPFC; Fig 4a) | 1 | R Frontal Medial Cortex<br>L Frontal Medial Cortex<br>R Frontal Pole | 24<br>23<br>10 | 58 | 2 | 56 | -18 | 3.21 |
| High-value Go, modulation across items (Fig 4b) | 1 | L Frontal Pole | 140 | 147 | -28 | 46 | -15.5 | 3.59 |
| High-value Go, modulation across items, small volume correction (hippocampus; Fig 4c) | 1 | Right Hippocampus | 34 | 36 | 26 | -6 | -25.5 | 3.79 |
| High-value Go minus NoGo, negative correlation across participants, small volume correction (SPL; Fig 4d) | 1 | R Superior Parietal Lobule | 87 | 87 | 30 | -40 | 44.5 | 3.92 |

***Supplementary Table 3*. *Passive viewing task, one month after minus before CAT:*** *Regions with significant activation for the imaging contrasts of* ***Figure 4****. For each cluster, the list presents all regions from the Harvard-Oxford atlas that contained at least 10 active voxels within the cluster, as well as the X/Y/Z location for the peak activation in MNI space.*

| Contrast | Cluster | Region | Number of voxels in region | Cluster size | X | Y | Z | Peak z stat |
|---|---|---|---|---|---|---|---|---|
| Choices of high-value Go minus choices of high-value NoGo items (Fig 5a) | 1 | R Occipital Fusiform Gyrus | 60 | 131 | 34 | -60 | -18 | 3.86 |
| | | R Temporal Occipital Fusiform Cortex | 40 | | | | | |
| | | R Lingual Gyrus | 29 | | | | | |
| | 2 | L Central Opercular Cortex | 59 | 136 | -68 | -12 | 12 | 3.74 |
| | | L Postcentral Gyrus | 10 | | | | | |
| | 3 | R Central Opercular Cortex | 101 | 152 | 46 | -10 | 7 | 3.91 |
| | | R Heschl's Gyrus (includes H1 and H2) | 14 | | | | | |
| | | R Insular Cortex | 13 | | | | | |
| | 4 | L Middle Temporal Gyrus, temporooccipital part | 65 | 188 | -48 | -60 | 12 | 4.04 |
| | | L Lateral Occipital Cortex, inferior division | 44 | | | | | |
| | | L Angular Gyrus | 24 | | | | | |
| | | L Lateral Occipital Cortex, superior division | 22 | | | | | |
| | | L Supramarginal Gyrus, posterior division | 13 | | | | | |
| Choices of high-value Go minus choices of high-value NoGo items, negative correlation across participants (Fig 5b) | 1 | Right Thalamus | 103 | 137 | 8 | -26 | 2 | 3.91 |
| | 2 | Left Caudate | 70 | 148 | -10 | 0 | 7 | 4 |
| | | Left Thalamus | 17 | | | | | |
| | | Left Accumbens | 16 | | | | | |
| | 3 | R Frontal Operculum Cortex | 93 | 187 | 30 | 28 | 4.5 | 4.17 |
| | | R Insular Cortex | 22 | | | | | |
| | | R Frontal Orbital Cortex | 20 | | | | | |
| | | R Inferior Frontal Gyrus, pars triangularis | 12 | | | | | |
| | 4 | L Frontal Operculum Cortex | 76 | 269 | -44 | 22 | 12 | 4.51 |
| | | L Frontal Orbital Cortex | 52 | | | | | |
| | | L Inferior Frontal Gyrus, pars opercularis | 31 | | | | | |

| | | | | | | | | |
|---|---|---|---|---|---|---|---|---|
| | | L Insular Cortex | 21 | | | | | |
| | 5 | R Superior Frontal Gyrus | 152 | 404 | 2 | 12 | 52 | 4.28 |
| | | L Juxtapositional Lobule Cortex (formerly Supplementary Motor Cortex) | 77 | | | | | |
| | | R Juxtapositional Lobule Cortex (formerly Supplementary Motor Cortex) | 60 | | | | | |
| | | R Paracingulate Gyrus | 33 | | | | | |
| | | L Superior Frontal Gyrus | 15 | | | | | |
| Choices of high-value Go items, negative modulation across items (Fig 5c) | 1 | R Superior Frontal Gyrus | 99 | 143 | 26 | 8 | 60 | 3.98 |
| | | R Middle Frontal Gyrus | 23 | | | | | |
| | 2 | L Central Opercular Cortex | 66 | 158 | -48 | -18 | 17 | 3.86 |
| | | L Parietal Operculum Cortex | 62 | | | | | |
| | | L Planum Temporale | 20 | | | | | |
| | 3 | L Frontal Pole | 93 | 225 | -40 | 30 | 35 | 4.51 |
| | | L Middle Frontal Gyrus | 86 | | | | | |
| | 4 | R Paracingulate Gyrus | 150 | 242 | 8 | 12 | 27 | 3.65 |
| | | R Cingulate Gyrus, anterior division | 42 | | | | | |
| | | R Juxtapositional Lobule Cortex (formerly Supplementary Motor Cortex) | 25 | | | | | |
| | 5 | Right Putamen | 58 | 245 | 32 | 16 | 2 | 3.88 |
| | | R Insular Cortex | 47 | | | | | |
| | | Right Caudate | 29 | | | | | |
| | | R Frontal Orbital Cortex | 23 | | | | | |
| | | R Frontal Operculum Cortex | 16 | | | | | |
| | 6 | L Insular Cortex | | 280 | -40 | 8 | 9.5 | 4.43 |
| | | L Frontal Operculum Cortex | 57 | | | | | |
| | | L Central Opercular Cortex | 47 | | | | | |
| | | Left Putamen | 27 | | | | | |
| | | L Frontal Orbital Cortex | 22 | | | | | |
| | 7 | L Precentral Gyrus | 172 | 365 | -26 | -6 | 67 | 4.15 |
| | | L Superior Frontal Gyrus | 90 | | | | | |

| | | | | | | | | |
|---|---|---|---|---|---|---|---|---|
| | | L Middle Frontal Gyrus | 30 | | | | | |
| Choices of high-value Go items, negative correlation across participants, small volume correction (SPL; Fig 5d) | 1 | R Superior Parietal Lobule | 53 | 53 | 34 | -48 | 47 | 3.44 |
| Choices of high-value Go items, negative modulation across items, small volume correction (SPL; Fig 5e) | 1 | R Superior Parietal Lobule | 47 | 47 | 26 | -44 | 67 | 3.8 |

***Supplementary Table 4. Probe task, after CAT:*** *Regions with significant activation for the imaging contrasts of* ***Figure 5a-e****. For each cluster, the list presents all regions from the Harvard-Oxford atlas that contained at least 10 active voxels within the cluster, as well as the X/Y/Z location for the peak activation in MNI space.*

| Contrast | Cluster | Region | Number of voxels in region | Cluster size | X | Y | Z | Peak z stat |
|---|---|---|---|---|---|---|---|---|
| Choices of high-value Go items, modulation across items (Fig 5f) | 1 | R Precuneous Cortex | 117 | 151 | 6 | -70 | 44.5 | 3.67 |
| | | L Precuneous Cortex | 30 | | | | | |
| | 2 | R Middle Temporal Gyrus, posterior division | 92 | 175 | 54 | -18 | -3 | 3.89 |
| | | R Superior Temporal Gyrus, posterior division | 62 | | | | | |
| | | R Superior Temporal Gyrus, anterior division | 14 | | | | | |
| | 3 | L Lateral Occipital Cortex, superior division | 108 | 201 | -46 | -64 | 24.5 | 4.18 |
| | | L Angular Gyrus | 62 | | | | | |
| | 4 | R Lateral Occipital Cortex, superior division | 205 | 208 | 52 | -66 | 34.5 | 3.87 |
| | 5 | L Middle Temporal Gyrus, posterior division | 226 | 342 | -56 | -24 | -10.5 | 4.3 |
| | | L Superior Temporal Gyrus, posterior division | 69 | | | | | |
| | | L Middle Temporal Gyrus, temporooccipital part | 23 | | | | | |
| Choices of high-value Go items, correlation across participants (Fig 5g) | 1 | R Postcentral Gyrus | 68 | 181 | 64 | -14 | 22 | 4.4 |
| | | R Parietal Operculum Cortex | 42 | | | | | |
| | | R Planum Temporale | 19 | | | | | |
| | | R Supramarginal Gyrus, anterior division | 18 | | | | | |
| | 2 | L Precuneous Cortex | 134 | 474 | 2 | -42 | 57 | 3.8 |
| | | R Precuneous Cortex | 121 | | | | | |
| | | R Cingulate Gyrus, posterior division | 111 | | | | | |
| | | L Cingulate Gyrus, posterior division | 81 | | | | | |
| | | L Intracalcarine Cortex | 10 | | | | | |

***Supplementary Table 5. Probe task, one month after CAT:*** *Regions with significant activation for the imaging contrasts of* ***Figure 5f-g****. For each cluster, the list presents all regions from the Harvard-Oxford atlas that contained at least 10 active voxels within the cluster, as well as the X/Y/Z location for the peak activation in MNI space.*
